# Numerical aspect of Predator-Pray Model

**DOI:** 10.1101/065516

**Authors:** Monika Justyna Nir, Sarangam Majumdar

## Abstract

In a closed eco-system, there are only two types of animals: the predator and the prey. They form a simple food-chain where the predator species hunts the prey species, while the prey grazes vegetation. The size of the two populations can be described by a simple system of two nonlinear first order differential equations formally known as the Lotka-Volterra equations, which originated in the study of fish populations of the Mediterranean during and immediately after World War I. Here, we study numerically this nonlinear parabolic evolution problem and compare the result of various numerical schemes.

## 1 Introduction

When species interact the population dynamics of each species is affected. In general there is a whole web of interacting species, sometimes called a trophic web, which makes for structurally complex communities (Abrams et al., 2000)(Aziz-Alaoui et al., 2003). We consider here systems involving 2 or more species, concentrating particularly on two-species systems.

There are three main types of interaction.

1. The growth rate of one population is decreased and the other increased the populations are in a predator-prey situation.
2. If the growth rate of each population is decreased then it is competition.
3. If each populations growth rate is enhanced then it is called mutualism or symbiosis (Murray, 2002).

Last few decades, thousand’s of papers are published in predation-pray model to investigate the dynamics of the system. The stage-structured predator-prey model and optimal harvesting policy is studied by (Zhang et al., 2000). Faria investigate stability and bifurcation for a delayed predator-prey model and the effect of diffusion (Faria et al., 2001). From the numerical point of view (Ahmed et al., 2007) find the numerical solutions of fractional order predator-prey and rabies models and Garvie, Marcus R. studied Finite-difference schemes for reaction-diffusion equations modeling predator-prey interactions in matlab (Garvie, 2007). A non-standard numerical scheme for a generalized Gause-type predator-prey model is also introduced by Moghadas, S. M., M. E. Alexander, and B. D. Corbett (Moghadas et al., 2004). Bandyopadhyay, M., and J. Chattopadhyay studied Ratio-dependent predator-prey model with the effect of environmental fluctuation and stability (Bandyopadhyay and Chattopadhyay, 2005). Pascual, Mercedes presented the diffusion-induced chaos in a spatial predator-prey system (Pascual, 1993). In this paper, we investigate some stiff and non-stiff numerical scheme and compare them for predator-pray model.

## 2 Mathematical Model

Volterra (1926) first proposed a simple model for the predation of one species by another to explain the oscillatory levels of certain fish catches in the Adriatic. If *N*(*t*) is the prey population and *P*(*t*) that of the predator at time t then Volterra’s model is

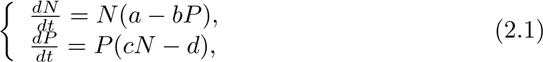

where *a, b, c* and *d* are positive constants.

The assumptions in the model are:

- The prey in the absence of any predation grows unboundedly in a Malthusian way; this is the*aN* term in the first equation of (1).
- The effect of the predation is to reduce the prey’s per capita growth rate by a term proportional to the prey and predator populations; this is the *−bNP* term.
- In the absence of any prey for sustenance the predator’s death rate results in exponential decay, that is, the *−dP* term in the second equation of (1).
- The prey’s contribution to the predators’ growth rate is *cNP*; that is, it is proportional to the available prey as well as to the size of the predator population.

The *NP* terms can be thought of as representing the conversion of energy from one source to another: *bNP* is taken from the prey and *cNP* accrues to the predators. We shall see that this model has serious drawbacks. Nevertheless it has been of considerable value in posing highly relevant questions and is a jumping *−*off place for more realistic models. The model (1) is known as the Lotka-Volterra model.

## 3 Numerical Simulations

In this section, we are studied the model using numerical explicit and implicit scheme and reported their behaviouras follows.

**Figure 1:**
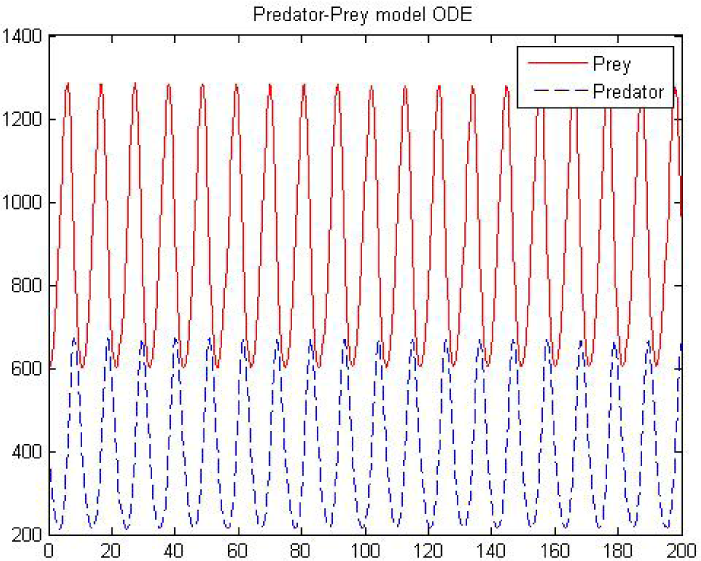
Predator- pray behaviour by ode23

**Figure 2:**
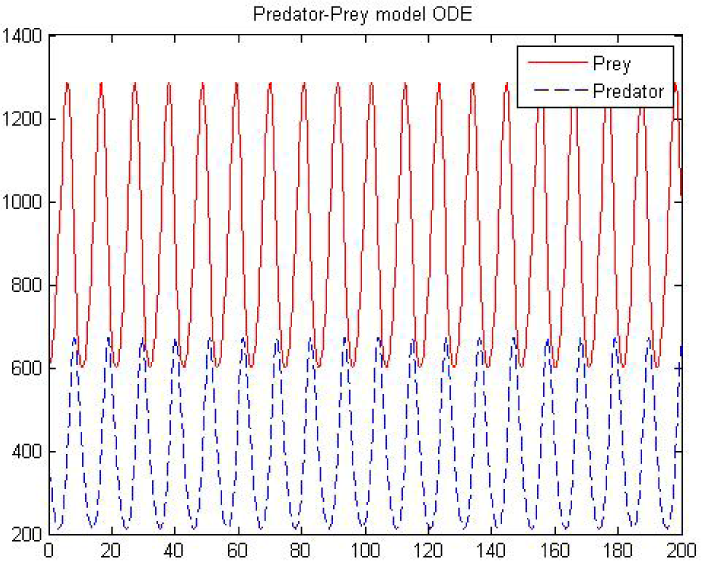
Predator- pray behaviour by ode45

**Figure 3:**
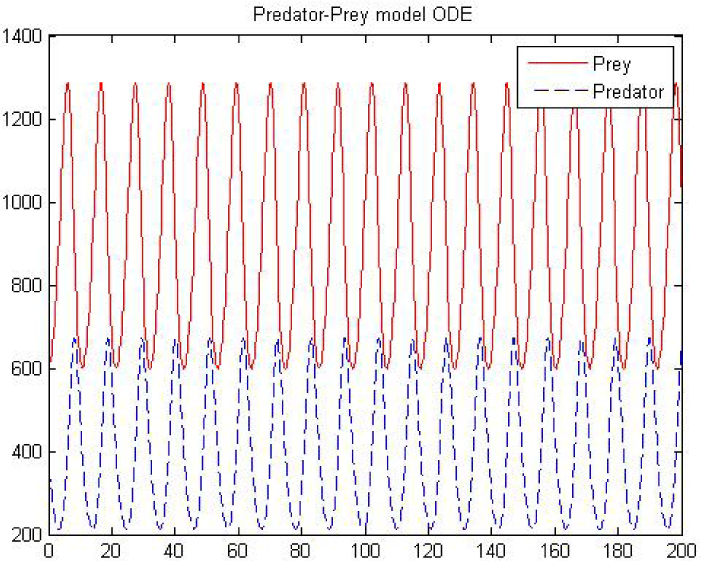
Predator- pray behaviour by ode23s

**Figure 4:**
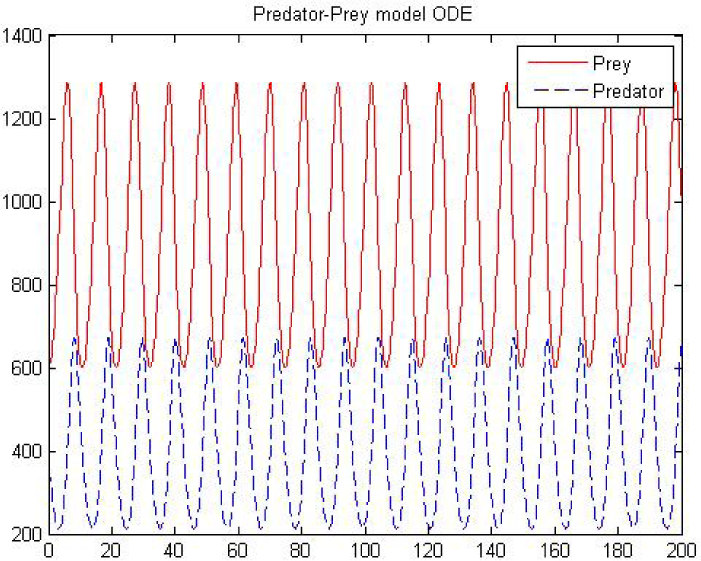
Predator- pray behaviour by ode23t

**Figure 5:**
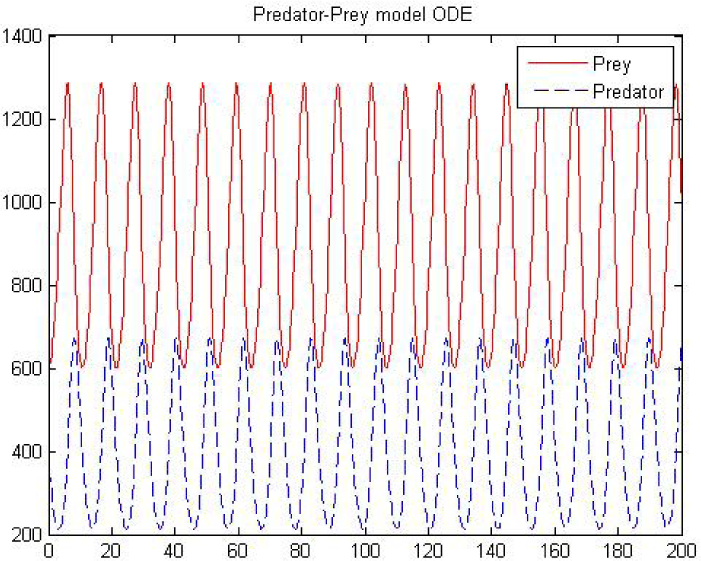
Predator- pray behaviour by ode23tb

**Figure 6:**
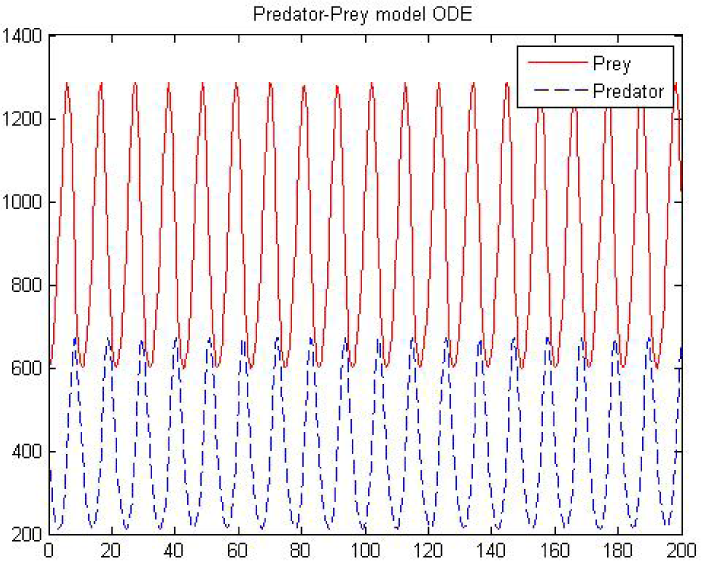
Predator- pray behaviour by ode113

**Table 1:** List of ode solver

| Function | Description |
| --- | --- |
| ode45 | Nonstiff differential equations, medium order method. |
| ode23 | Nonstiff differential equations, low order method. |
| ode113 | Nonstiff differential equations, variable order method. |
| ode15s | Stiff differential equations and DAEs, variable order method. |
| ode23s | Stiff differential equations, low order method. |
| ode23t | Moderately stiff differential equations and DAEs, trapezoidal rule. |
| ode23tb | Stiff differential equations, low order method. |

**Figure 7:**
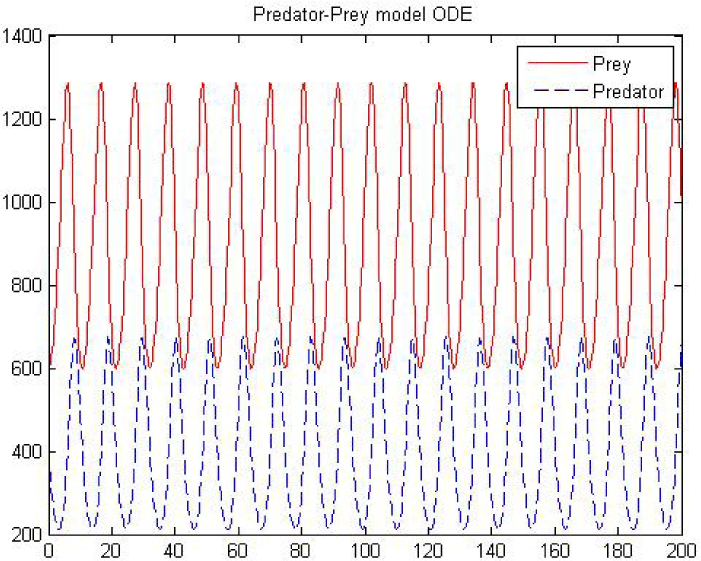
Predator- pray behaviour by ode15s

We used ode23tb, ode23s, ode15s, ode23t which are known as stiff solver. For a stiff problem, solutions can change on a time scale that is very short compared to the interval of integration, but the solution of interest changes on a much longer time scale. Methods not designed for stiff problems are ineffective on intervals where the solution changes slowly because they use time steps small enough to resolve the fastest possible change. Stiff solvers can be used exactly like the other solvers. However, we can often significantly improve the efficiency of the stiff solvers by providing them with additional information about the problem. We use

1. ode23tb which is an implementation of TR-BDF2, an implicit Runge-Kutta formula with a first stage that is a trapezoidal rule step and a second stage that is a backward differentiation formula of order two. By construction, the same iteration matrix is used in evaluating both stages. Figure 5 shows the behaviour of predator- pray model.

2. we use ode23t solver. ode23t is an implementation of the trapezoidal rule using a free interpolant. If the problem is only moderately stiff and we need a solution without numerical damping, then we usually use this solver. Figure 4, demonstrate predator-pray behaviour in three dimension using ode23t at different numbers of grids.

3.ode23s is based on a modified Rosenbrock formula of order 2. Because it is a one-step solver, it may be more efficient than ode15s at crude tolerances. It can solve some types of stiff problems for which ode15s is not effective. Figure 3 show the behaviour of the predator-pray using ode23s when we are increasing number of nodes.

4. Finally, we choose ode15s solver. ode15s is a variable-order solver based on the numerical differentiation formulas (NDFs). Optionally it uses the backward differentiation formulas, BDFs, (also known as Gears method) that are usually less efficient. Like ode113, ode15s is a multistep solver. If we suspect that a problem is stiff or if ode45 failed or was very inefficient, then ode15s is the ultimate choice. In this solver, the maximum order is an integer 1 through 5 used to set an upper bound on the order of the formula that computes the solution. By default, the maximum order is 5. Moreover, set BDF on to have ode15s use the BDFs. By default, BDF is off, and the solver uses the NDFs. For both the NDFs and BDFs, the formulas of orders 1 and 2 are A-stable (the stability region includes the entire left half complex plane). The higher order formulas are not as stable, and the higher the order the worse the stability. Figure 7, illustrate predator-pray behaviour in the spatio temporal complex phenomenon using ode15s.

The performance of the stiff solvers varies depending on the format of the problem and specified options, provided the Jacobian matrix or sparsity pattern always improves solver efficiency for stiff problems. But since the stiff solvers use the Jacobian differently, the improvement can vary significantly. Practically speaking, if a system of equations is very large or needs to be solved many times, then it is worthwhile to investigate the performance of the different solvers to minimize execution time. In the case of the Lotka-Voltera equation, ode23s works efficiently in less amount of time. Other solvers are also able to solve the L-V equation with longer times than ode23s. From the point of view of absolute error ode15s is a better solver than other in this case. If we compare with stiff and non-stiff solver. Then we can say that stiff solver take less time than non-stiff.

**Figure 8:**
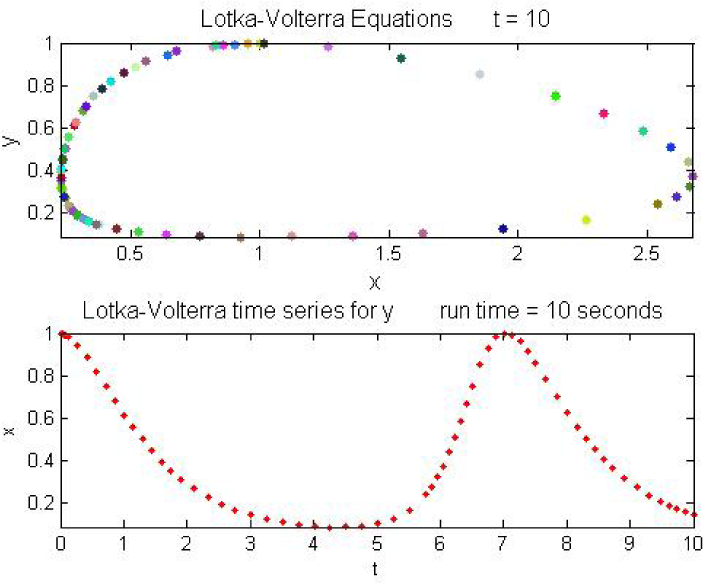
Phase plane and time series

**Figure 9:**
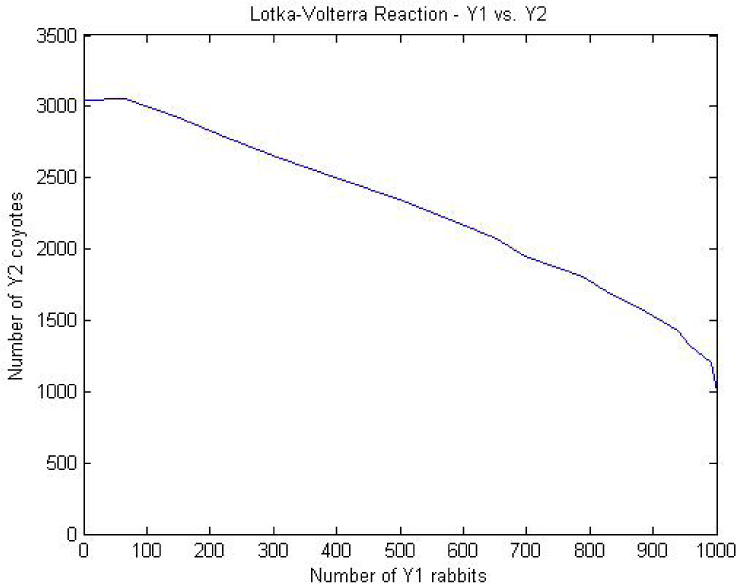
Competition between two species

Figure 8 illustrate phase plane and the time series for *a* = 1, *b* = 2.666667, *c* = 1 and *d* = 1 Moreover, Figure 9 shows the competition between two species.

## 4 Conclusion

We use different numerical scheme for the predator-pray model. We observed that the stiff solver are more efficient than than the non-stiff solver. We conclude that the stiffness is a subtle, difficult, and important concept in the numerical solution of ordinary differential equations. It depends on the differential equation, the initial conditions, and the numerical method. Dictionary definitions of the word “stiff” involve terms like “not easily bent”, “rigid”, and “stubborn”. We are concerned with a computational version of these properties. An ordinary differential equation problem is stiff if the solution being sought is varying slowly, but there are nearby solutions that vary rapidly, so the numerical method must take small steps to obtain satisfactory results. Stiffness is an efficiency issue. If we weren’t concerned with how much time a computation takes, we wouldn’t be concerned about stiffness. Nonstiff methods can solve stiff problems; they just take a long time to do it. Nonetheless, we believe this example (Predator-Pray) is fairly representative with respect to many features.

